# Extracellular Electron Uptake by Two *Methanosarcina* Species

**DOI:** 10.1101/458091

**Authors:** Mon Oo Yee, Oona Snoeyenbos-West, Bo Thamdrup, Lars DM Ottosen, Amelia-Elena Rotaru

**Affiliations:** Nordcee, Department of Biology, University of Southern Denmark, Odense, Denmark; Deparment of Engineering, University of Aarhus, Aarhus, Denmark

## Abstract

Direct electron uptake by prokaryotes is a recently described mechanism with a potential application for energy and CO_2_ storage into value added chemicals. Members of Methanosarcinales, an environmentally and biotechnologically relevant group of methanogens, were previously shown to retrieve electrons from an extracellular electrogenic partner performing Direct Interspecies Electron Transfer (DIET) and were therefore proposed to be electroactive. However, their intrinsic electroactivity has never been examined. In this study, we tested two methanogens belonging to the genus *Methanosarcina, M. barkeri* and *M. horonobensis,* regarding their ability to accept electrons directly from insoluble electron donors like other cells, conductive particles and electrodes. Both methanogens were able to retrieve electrons from *Geobacter metallireducens* via DIET. Furthermore, DIET was also stimulated upon addition of electrically conductive granular activated carbon (GAC) when each was co-cultured with *G. metallireducens*. However, when provided with a cathode poised at −400 mV (vs. SHE), only *M. barkeri* could perform electromethanogenesis. In contrast, the strict hydrogenotrophic methanogen, *Methanobacterium formicicum*, did not produce methane regardless of the type of insoluble electron donor provided (*Geobacter* cells, GAC or electrodes). A comparison of functional gene categories between the two *Methanosarcina* showed differences regarding energy metabolism, which could explain dissimilarities concerning electromethanogenesis at fixed potentials. We suggest that these dissimilarities are minimized in the presence of an electrogenic DIET partner (e.g. *Geobacter*), which can modulate its surface redox potentials by adjusting the expression of electroactive surface proteins.

## Introduction

Extracellular electron uptake by methanogens may impact carbon turnover in electron-acceptor limited environments (Morris et al., 2013). In these environments, thermodynamically challenging processes become possible due to syntrophic interactions between bacteria and archaea. A syntrophic interaction requires a bacterium, which oxidizes organics to interspecies-transferrable molecules. Moreover, syntrophy requires a partner methanogen to scavenge the transferable molecules. For decades, we have assumed that interspecies-transferrable molecules were either H2 or formate (Stams and Plugge, 2009). We now know that some species can also transfer electrons directly (Lovley, 2017). During direct interspecies electron transfer (DIET), species like *Geobacter* oxidize ethanol according to reaction (1), only in the presence of methanogens like *Methanosarcina*, which scavenge reducing equivalents (H^+^and e^−^) and acetate (Rotaru et al., 2014a, 2014b) (Fig. 1).

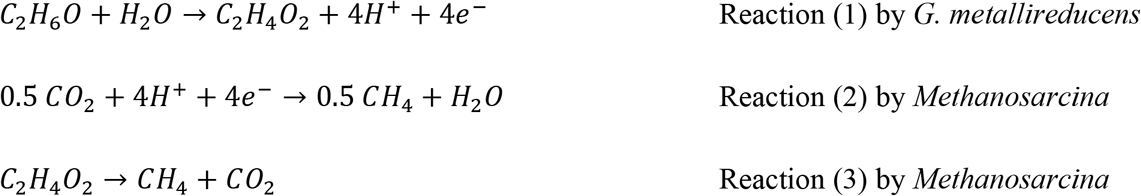

**Fig. 1.**
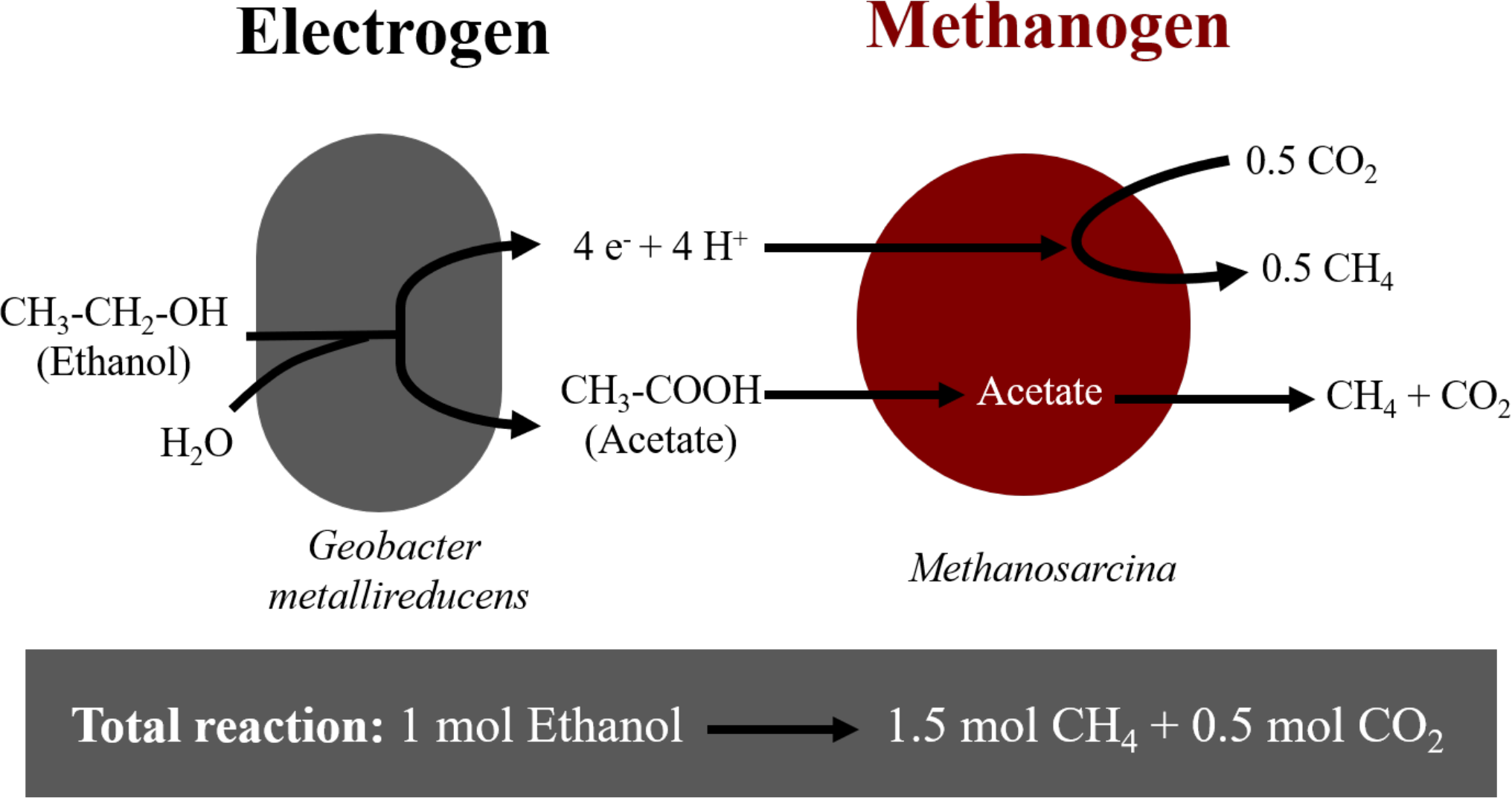
Diagram of electron flow during DIET interactions between *Geobacter metallireducens* and *Methanosarcina*-species provided with ethanol as sole electron donor.

In DIET co-cultures, only those *Geobacter* species producing high current densities, met the energetic needs of their DIET partners (Rotaru et al., 2015). For this purpose, *Geobacter* up-regulates the expression of redox active and conductive proteins (outer membrane *c*-type cytochromes and pili) (Holmes et al., 2018b; Shrestha et al., 2013). *Geobacter*’s requirement for outer-surface proteins during DIET was confirmed earlier with gene-deletion studies (Rotaru et al., 2014a, 2014b). Thus, if *Geobacter* lacked the ability to produce e.g. pili it was incapable of DIET.

Although we understand how *Geobacter* releases electrons outside their cells during DIET, the way Methanosarcinales retrieve DIET-electrons is poorly understood. A glimpse at this mechanism was provided in a recent comparative transcriptomic study (Holmes et al., 2018b). In this study, the transcriptomes of DIET co-cultures (*G. metallireducens – Methanosarcina barkeri*) were compared to those of co-cultures performing interspecies H2-transfer (*Pelobacter carbinolicus – M. barkeri*). During DIET, *M. barkeri* had higher expression of membrane-bound redox-active proteins like cupredoxins, thioredoxins, pyrroloquinoline, and quinone-, cytochrome-or Fe-S containing proteins (Holmes et al., 2018b). Still, the exact mechanism of electron uptake by *Methanosarcina* has not been validated and warrants further investigation.

Moreover, *Methanosarcina* can also retrieve electrons from electrically conductive particles charged by *Geobacter* oxidizing organics (Shrestha and Rotaru, 2014). In effect, DIET is accelerated by electrically conductive particles/minerals perhaps because they replace conductive and redox active surface proteins diminishing cellular energy expenditure required to overexpress such surface constituents (Liu et al., 2012). For instance, co-cultures of *G. metallireducens* and *M. barkeri* were stimulated at the addition of conductive particles, such as GAC (Liu et al., 2012), carbon cloth (Chen et al., 2014a), biochar (Chen et al., 2014b), or magnetite (Wang et al., 2018). On the other hand, the addition of non-conductive materials did not stimulate DIET (Chen et al., 2014a; Rotaru et al., 2018a). In addition, conductive particles appear to play a significant role in interspecies interactions from natural and artificial ecosystems such as sediments, soils, rice paddies or anaerobic digesters (Holmes et al., 2017a; Rotaru et al., 2018b; Ye et al., 2018; Zhang et al., 2018). In these cases, the addition of conductive particles enriched for DIET-related Methanosarcinales (Dang et al., 2016; Holmes et al., 2017b; Rotaru et al., 2018a; Zheng et al., 2015). However, exceptions were observed since occasionally conductive particles enriched for H2-utilizing methanogens of the genus *Methanospirillum* (Lee et al., 2016) or *Methanobacterium* (Lin et al., 2017; Zhuang et al., 2015).

Since methanogens retrieve extracellular electrons from cells or conductive particles to reduce CO_2_ to methane, it was expected that they could also retrieve electrons from a poised electrode via electromethanogenesis. Nevertheless, electromethanogenesis was only verified with H_2_-utilizing methanogens like *Methanobacterium palustre* (Cheng et al., 2009). Yet, H_2_-utilizers were incapable of DIET (Rotaru et al., 2014b). Conversely, it is unknown if Methanosarcinales, which are capable of DIET, are also capable of electron uptake from a cathode. Our objective was to investigate the ability to carry electromethanogenesis and DIET in two *Methanosarcina* species. We have shown that both *Methanosarcina* species grew by DIET, however only *M. barkeri* grew on the cathode at − 400 mV vs. SHE. This indicates that extracellular electron-uptake routes from cathodes or other cells might differ between *Methanosarcina* species.

## Materials and Methods

### Microorganism strains and cultivation conditions

*Methanosarcina barkeri* MS (DSM 800) and *Methanosarcina horonobensis* HB-1 (DSM 21571) were purchased from the German culture collection DSMZ while *Methanobacterium formicicum* (NBRC 100475) was from the Japanese culture collection NBRC. *Geobacter metallireducens* GS-15 was sent to us by Dr. Sabrina Beckmann from the University of New South Wales, Australia.

Routine cultivation was performed under strictly anaerobic conditions in serum bottles sealed with butyl rubber stoppers and incubated statically at 37°C. All the microorganisms had been adapted to grow in DSMZ medium 120c with the following modifications: 1 g/L NaCl, 0.5g/L yeast, and no tryptone (Rotaru et al., 2014a). During incubations in co-cultures or for electrochemical experiments, sulfide and yeast extract was omitted. When grown in pure cultures, *Methanosarcina* species were provided with 30 mM acetate and 20 mM methanol as methanogenic substrates, while *M. formicicum* was provided with 150 kPa of H_2_: CO_2_ (80:20) in the headspace. *G. metallireducens* was routinely grown with 20 mM ethanol and 56 mM ferric citrate. All media and cultures were prepared and kept under an N_2_: CO_2_ (80:20) atmosphere.

The co-cultures of *Geobacter* and methanogens were initiated with 0.5 mL of *G. metallireducens* and 1 mL of acetate-grown *Methanosarcina-*species or H_2_-grown *M. formicicum* inoculated into 8.5 ml of the media prepared as above. The starting cell numbers for the methanogens in co-cultures were approximately 2.6 × 10^6^ cells/mL, 8.2 × 10^6^ cells/mL and 6.7 × 10^5^ cells/mL for *M. barkeri, M. formicicum* and *M. horonobensis*, respectively. The starting cell numbers for *G. metallireducens* in co-cultures were circa 1.5 × 10^5^. Incubations were carried out in 20 ml pressure vials. For the co-cultures ethanol (10 mM) was added as an electron donor and CO_2_ was the sole electron acceptor. When noted, sterile granular activated carbon (GAC) was added at a concentration of 25 g/L and prepared as described before (Rotaru et al., 2018a).

### Electrochemical setup and measurements

All bioelectrochemical incubations were carried with a modified DSMZ 120c medium (see above) in the absence of sulfide and yeast extract. The pH of this medium in the bioelectrochemical reactors was set to 6.5. We used bioelectrochemical reactors with a standard dual chamber configuration as shown in **Figure S1**. Two-chamber glass bottles were purchased from Adams & Chittenden Scientific Glass (USA) with side ports fitted with butyl septa to allow for medium transfer, sampling, and the introduction of a reference electrode. Each chamber of the reactors had a total volume of 650 ml with a flange diameter of 40 mm and the chambers were separated by a Nafion™ N117 proton exchange membrane (Ion Power) held by an O-ring seal with a knuckle clamp.

Both the working and counter electrodes were made of graphite (Mersen MI Corp., Greenville USA) with dimensions of 2.5 cm × 7.5 cm × 1.2 cm thus a total projected surface area of 61.5 cm^2^. The working and counter electrodes were coupled to titanium wires, which pierced through rubber stoppers fitted into the main opening of each chamber. A 2 cm deep and 2 mm wide hole was drilled on the short side of the electrode and a 12.5 cm long; 2 mm wide titanium rod (Alfa-Aesar, DE) was inserted and sealed from the outside with biocompatible non-conductive epoxy. Electrodes with a resistance of less than 10 Ω were used to ensure proper electrical connections. The assembled electrodes were introduced into the chamber via the main opening and 2 mm-wide holes were drilled in the black rubber stopper to allow access of the titanium rod. After autoclaving the reactors, sterile medium was transferred into the reactors anaerobically and aseptically. Sterile (bleach and ethanol series) reference electrodes were lodged through a side port in the working electrode chamber at a distance of about 1 cm from the surface of the working electrode. After lodging the electrodes, degassing with N_2_: CO_2_ (80:20) for circa 30 minutes in each reactor chamber ensured anaerobic conditions. When the pre-cultures were in mid-exponential phase, they were inoculated (20%) into fresh medium in the cathodic chamber following sterile anoxic techniques to a final volume of 550 ml leaving a headspace of approximately 100 mL in each chamber. The approximate cell numbers at the time of inoculation into the electrochemical reactors for *M. barkeri, M. formicicum* and *M. horonobensis* were 2.6 × 10^7^ cells/mL, 8.2 × 10^7^ cells/mL and 6.7 × 10^6^ cells/mL respectively. Cell counts were done with microscopic examination using DAPI (1 μg/mL) stained cells.

The reference electrodes used were leak-free Ag/AgCl reference electrodes (3.4M KCl) (CMA Microdialysis, Sweden), which are 242 mV above the standard hydrogen electrode (SHE) according to the manufacturer and our own measurements against a Hydroflex^®^ reference electrode used as NHE (normal hydrogen electrode). The difference between NHE and SHE is experimentally negligible (Smith and Stevenson, 2007). All potentials in this paper from here onwards are reported vs. SHE by adjusting accordingly from the Ag/AgCl reference electrodes values. Cathode poising and electrochemical measurements were carried with a multichannel potentiostat (MultiEmstat, Palmsens, NL) operated by the Multitrace software (Palmsens, NL).

### Analytical measurements and calculations

Headspace samples for CH4 and H_2_ analysis were taken with hypodermic needles and kept in airtight exetainers until measurement. Methane (CH_4_) and hydrogen gas (H_2_) were measured on a Trace 1300 gas chromatograph (Thermo-Scientific) with a TracePLOT™ TG-BOND Msieve 5A column and a thermal conductivity detector (TCD). The carrier gas was argon at a flow rate of 25 mL/min. The injector, oven and detector temperatures were 150°C, 70°C, and 200°C respectively. The detection limit for CH_4_ and H_2_ was ca. 5 μM for both. The concentration unit was converted to molarity by using the ideal gas law (p×V = n×R×T) under standard conditions, where p = 1 atm, V is the volume of the gaseous phase (L), n is amount of gas (mol), R is the gas constant (0.08205 atm×L/ mol×K) and T = 298.15 K. For ethanol detection, 0.5 mL samples were filtered (0.2 μm pore size) into appropriate sampling vials and were heated for 5 min. at 60°C. The headspace gas was then pass through the Trace 1300 gas chromatograph (Thermo-Scientific) with a TRACE™ TR-Wax column and detected by a flame ionization detector (FID). Nitrogen gas at a flow of 1 mL/min was used as the carrier and the injector, oven, and detectors were kept at 220°C, 40°C and 230°C respectively. Short-chained volatile fatty acids (VFA) were analyzed with a Dionex™ ICS-1500 Ion Chromatography system, using a Dionex™ IonPac™ AS22 IC Column and a mixture of 1.4 mM NaHCO_3_ and 4.5 mM Na_2_CO_3_ as the eluent fitted with an electron capture detector (ECD) at 30 mA.

### Genome comparison

Genomes for all tested microorganisms were available at the JGI integrated microbial genomes and microbiomes. Functional category comparisons and pairwise average nucleotide identity (ANI) were determined using the IMG/M-“Compare Genomes” tools. The IMG genome IDs of the studied *M. barkeri, M. horonobensis* and *M. formicicum* used were 2630968729, 2627854269 and 2645727909 respectively. The gene functions were analyzed from the annotated names of all the protein-coding genes retrieved from the National Center for Biotechnology Information (NCBI) database. The accession numbers used were NZ_CP009528, NZ_CP009516, and NZ_LN515531 for *M. barkeri, M. horonobensis* and *M. formicicum* respectively. To scan for the cytochrome motif (CxxCH) through all the genomes, we used a pattern-matching Web-application (Seiler et al., 2006) in addition to manual search

## Results and Discussion

It was previously shown that two Methanosarcinales-methanogens, *Methanosarcina barkeri* and *Methanothrix harundinacea* grew via DIET whereas strict H_2_-utilizing methanogens did not (Rotaru et al., 2014a, 2014b). Here we show that indeed *M. barkeri* could retrieve electrons not only from an exoelectrogen but also from an electrode poised at − 400 mV (non-H_2_ generating conditions) to carry electromethanogenesis. As expected, the H_2_-utilizing methanogen *M. formicicum* did not carry electromethanogenesis under this condition. We tested an environmentally relevant *Methanosarcina*, *M. horonobensis* for extracellular electron uptake from cells and electrodes, and we observed that it could only retrieve electrons from exoelectrogenic *Geobacter* and from granular activated carbon but not from electrodes.

### Methanosarcina barkeri

*barkeri* grows in co-culture with *G. metallireducens* via DIET, and the interaction could be accelerated by electrically conductive particles (Rotaru et al., 2014a, **Fig. S2**). This was anticipated because *G. metallireducens*, a respiratory organism, is incapable of substrate fermentation and consequent H_2_ production according to previous physiological tests (Cord-Ruwisch et al., 1998) and genetic investigations (Aklujkar et al., 2009). Since H_2_ could not be generated by *G. metallireducens*, a strict H_2_-utilizer like *M. formicicum* was rendered incapable of an interspecies association based on H_2_-transfer with this bacterium (Rotaru 2014b). However, more recent studies indicated that conductive carbon nanotubes stimulated methanogenesis by *M. formicicum* (Salvador et al., 2017). This implied that *M. formicicum* might be encouraged by the presence of conductive particle to interact syntrophically with *Geobacter*. Therefore, we tested if conductive GAC aids *M. formicicum* to establish a syntrophic association with *G. metallireducens*. This was not the case since co-cultures of *M. formicicum* and *G. metallireducens* did not generate methane regardless of the presence or absence of conductive particles, over the course of 120 days (Fig. 2).

**Fig. 2.**
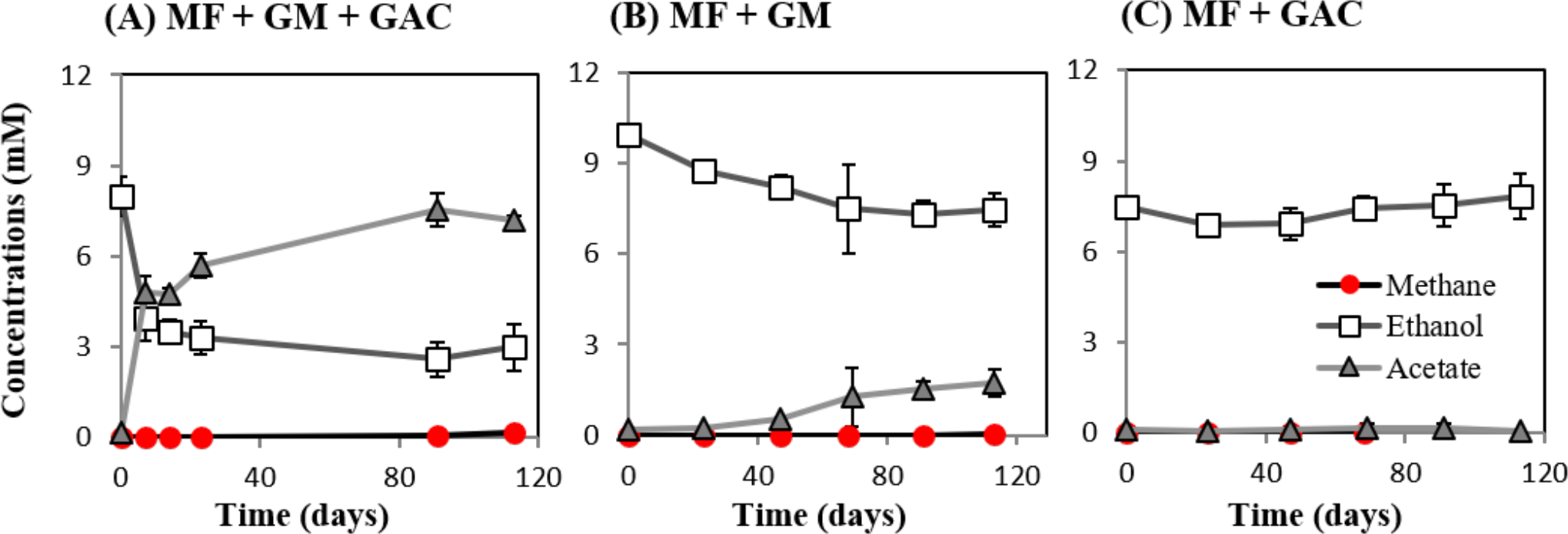
Co-cultures experiments with *M. formicicum* and *G.metallireducens* (n=3). *M. formicicum* did not produce methane when incubated with *G. metallireducens* in the presence (A) or absence of GAC (B). Alone, *M. formicicum* could not utilize ethanol or produce methane in the presence of GAC. MF - *Methanobacterium formicicum*, GM - *Geobacter metallireducens*, GAC - granular activated carbon.

During DIET, the extracellular electron transfer machinery of *G. metallireducens* plays a crucial role in *Geobacter-Methanosarcina* interactions, indicating that *Geobacter* releases extracellular electrons for the methanogen to use. Therefore, we suspected that *Methanosarcina* might also be able to directly retrieve extracellular electrons from electrodes to do electromethanogenesis.

In this study, we tested for the first time if *M. barkeri* could retrieve electrons directly from an electrode poised at – 400 mV. Indeed, *M. barkeri* produced significantly more methane (4.4 ± 0.33 mM; p<0.001) (Fig. 3) when provided with an applied potential at the cathode, in contrast to open circuit controls without an applied potential (1.3 ± 0.33 mM) **(Fig. S3)**. The background methane in control reactors resulted from carry-over substrates, once this was subtracted, the additional methane produced by *M. barkeri* in poised reactors (3.1 ± 0.34 mM) could be solely credited to electricity. Moreover, the highest rate of methane production was observed when current density profiles indicated the highest current draw by *M. barkeri* (Fig. 3).

**Fig. 3.**
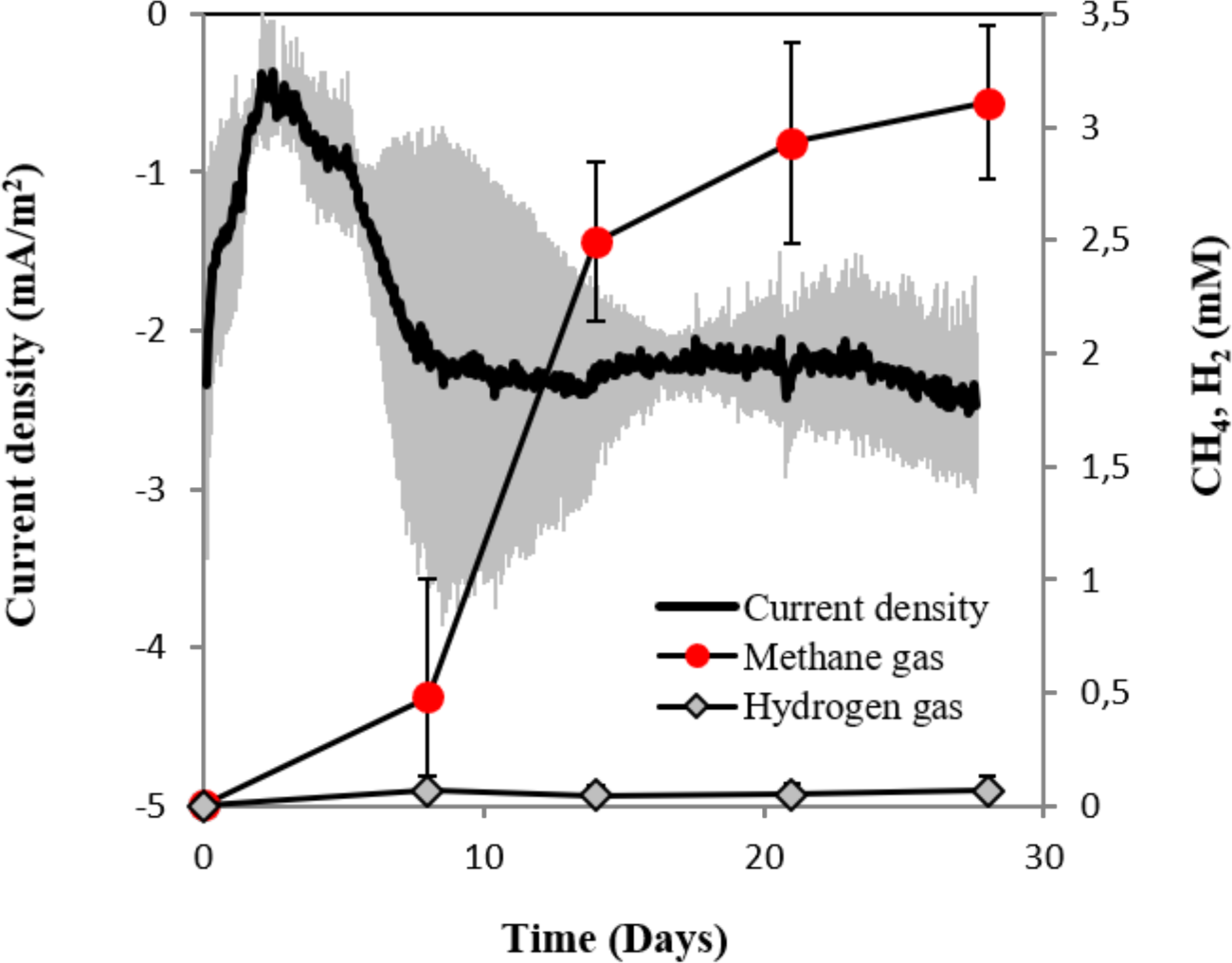
Current consumption and gas production in triplicate *M. barkeri* cultures provided with a poised cathode at − 400 mV (vs. SHE) as sole electron donor.

There are two possible scenarios for *M. barkeri* growing successfully at a cathode poised at – 400 mV:

1. It may use low concentrations of H_2_ generated electrochemically at the cathode, or
2. It retrieves electrons directly via an unknown mechanism

To determine abiotic electrochemical H_2_ evolution we i) verified for H_2_ accumulation over a month of incubation and ii) verified the threshold for H_2_-evolution by linear sweep voltammetry at the beginning and the end of the incubation. H_2_ did not accumulate over a month of incubation in six independent abiotic controls (Fig. 4A). Linear sweep voltammetry profiles indicated that in our media the threshold for H_2_-evolution was below –700 mV (Fig. 4B). This was in agreement with previous studies determining electrochemical H_2_-evolution under physiological conditions on a graphite electrode, which was below −400 mV due to high overpotentials (Batlle-Vilanova et al., 2014; Beese-Vasbender et al., 2015; Cheng et al., 2009; Mitov et al., 2012).

**Fig. 4.**
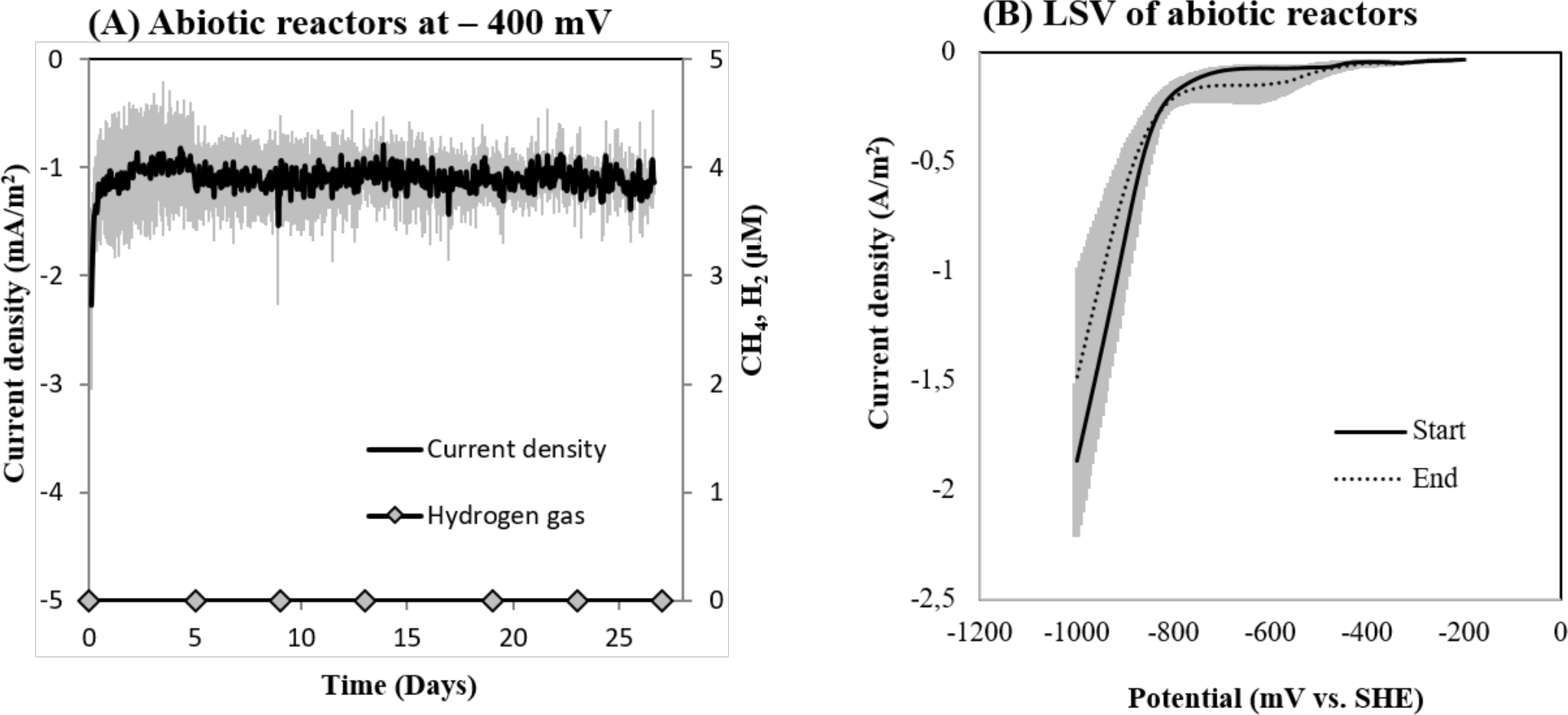
(A) Current consumption and gas production in four abiotic reactors at −400 mV (vs. SHE) and (B) Linear Sweep Voltammetry (LSV) of abiotic reactors at the start and end of the experiment (n=3).

On the other hand, in reactors inoculated with *M. barkeri*, the detected H_2_ stabilized at 0.065 ± 0.02 mM, similar to concentrations observed for co-cultures of *M. barkeri* with (0.077 ± 0.03 mM) or without conductive particles (0.076 ± 0.06 mM) and in pure culture (0.068 mM). This is supported by previous research, which demonstrated H_2_-cycling (H_2_-production and H_2_-uptake) in *M. barkeri* (Kulkarni et al., 2009, 2018; Mand et al., 2018). The cellular evolved H_2_ is well above the H_2_-uptake threshold for *M. barkeri* (296 nM −376 nM) (Kral et al., 1998; Lovley, 1985) possibly because in these cultures there is an alternative, competitive electron donor.

Secondly, if H_2_ evolved electrochemically to concentrations under the detection limit (which was not the case, see above), we anticipated that a sensitive hydrogenotrophic methanogen could effectively reclaim low concentrations of electrochemical H_2_, draw current and produce methane. To test this hypothesis we used a highly effective H_2_-utilizing methanogen – *M. formicicum*, which has a low H_2_ uptake threshold of approximately 6 nM (Lovley, 1985). However, when *M. formicicum* was incubated in electrochemical reactors, neither H_2_, methane nor current draw was observed at – 400 mV (Fig. 5) indicating that methanogenesis from H_2_ could not occur at this potential. In addition, to ensure that the growth of *M. formicicum* was unrestrained by the poised electrode, we carried control incubations at – 400 mV with extrinsic H_2_ as substrate. *M. formicicum* was unaffected by a poised electrode since it produced methane from the extrinsic H_2_ in an electrochemical setup **(Fig. S4)**.

**Fig. 5.**
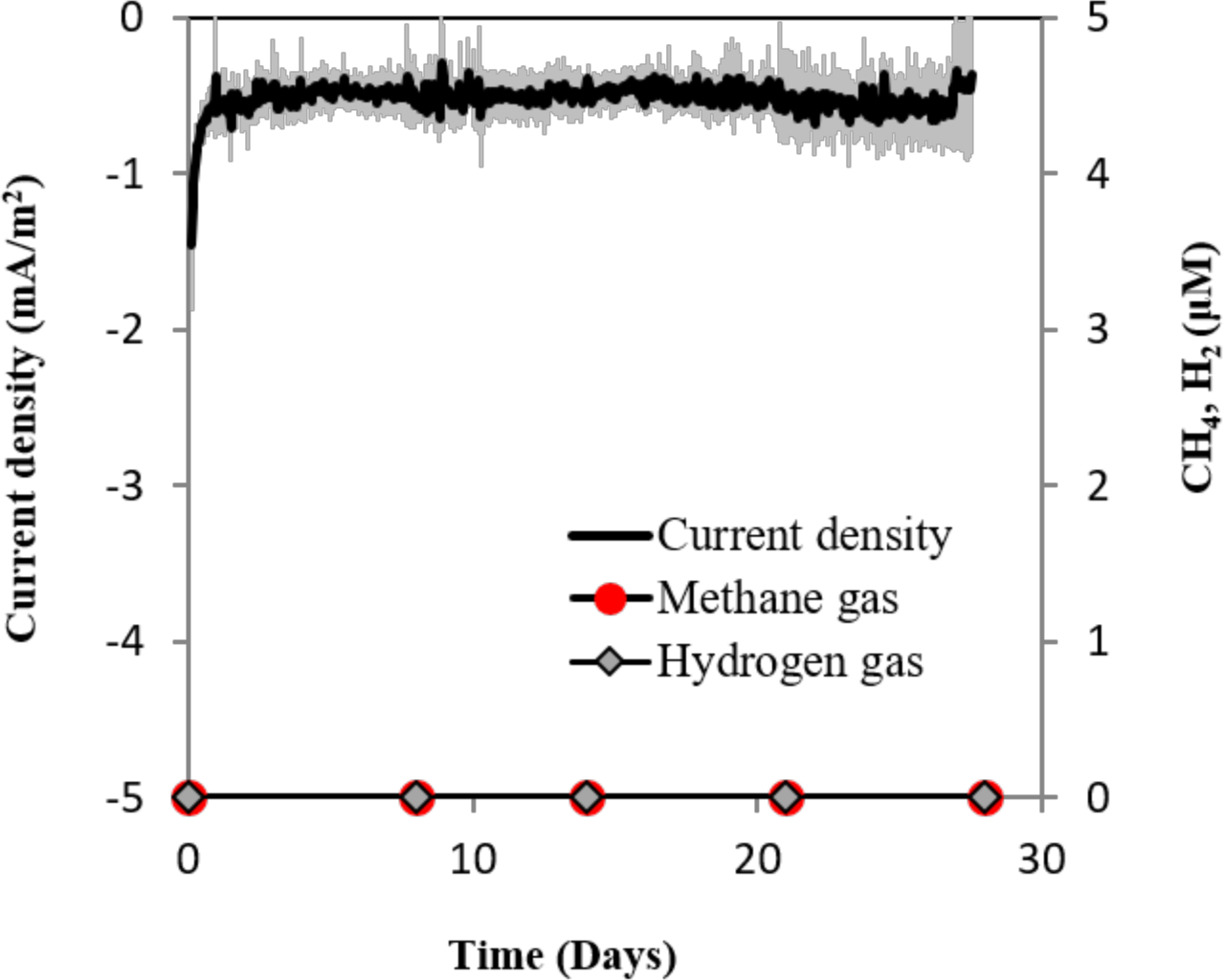
Current consumption and gas production in triplicate *M. formicicum* cultures provided with a cathode poised at − 400 mV (vs. SHE) as sole electron donor.

As electrochemical H_2_ was unlikely in our electrochemical setup, according to cumulative gas-detection analyses, electrochemical tests, and tests with a highly effective H_2_-utilizer, we confer that *M. barkeri* is likely to retrieve electrons directly from the electrode.

### Methanosarcina horonobensis

Except for *M. barkeri* and *M. harundinacea*, the distribution of extracellular electron uptake to other species of the order Methanosarcinales has not been explored. *M. barkeri* and *M. harundinacea* species have been isolated from and associated with anaerobic wastewater treatment (Bryant and Boone, 1987; De Vrieze et al., 2012; Ma et al., 2006). We were interested to see if other environmentally relevant *Methanosarcina* species had similar electron-uptake properties. We focused on *Methanosarcina horonobensis* because of its provenience and consistent association with deep aquifers (Holmes et al., 2018a; Shimizu et al., 2010).

*M. horonobensis* did establish successful syntrophic associations with *G. metallireducens* with or without conductive particles as an electrical conduit (Fig. 6). Theoretically, *G. metallireducens* oxidizes ethanol to acetate only if they could use the methanogen as an electron acceptor (Reaction 1). The acetate is then further disproportionated by the acetoclastic methanogen to produce methane and CO_2_ (Reaction 2 & 3). During DIET we expect the conversion of 1 mol ethanol to 1.5 mol methane according to Reactions 1 to 3 (Fig. 1). As predicted, in the *G. metallireducens – M. horonobensis* co-cultures, the syntroph oxidized 8.8 ± 0.4 mM ethanol providing the reducing equivalents (directly and via acetate) to generate 13.1 ± 0.8 mM CH_4_ by the methanogen. These co-cultures achieved stoichiometric recoveries of 98.5 ± 3.3 %. Similar recoveries (109 ± 18.5 %) were also observed at the addition of conductive particles. Single species controls with GAC showed that ethanol could not be converted to methane by the methanogen or the syntroph alone (Fig. 6C **& S5**). However, similar to previous reports (Zhang et al., 2018), *Geobacter* could partially convert ethanol to acetate using GAC as insoluble electron acceptor (Fig. S5; Van Der Zee et al., 2003; Zhang et al., 2018), likely until it reaches its maximum capacitance of 40 F/g (Zhang et al., 2009). Co-cultures of *G. metallireducens* and *M. horonobensis* could not carry interspecies H_2_ transfer because *G. metallireducens* is a strict respiratory microorganism which cannot ferment ethanol to generate H_2_ (Shrestha et al., 2013) and because *M. horonobensis* is unable to use H_2_ as electron donor for their metabolism (Shimizu et al., 2010).

**Fig. 6.**
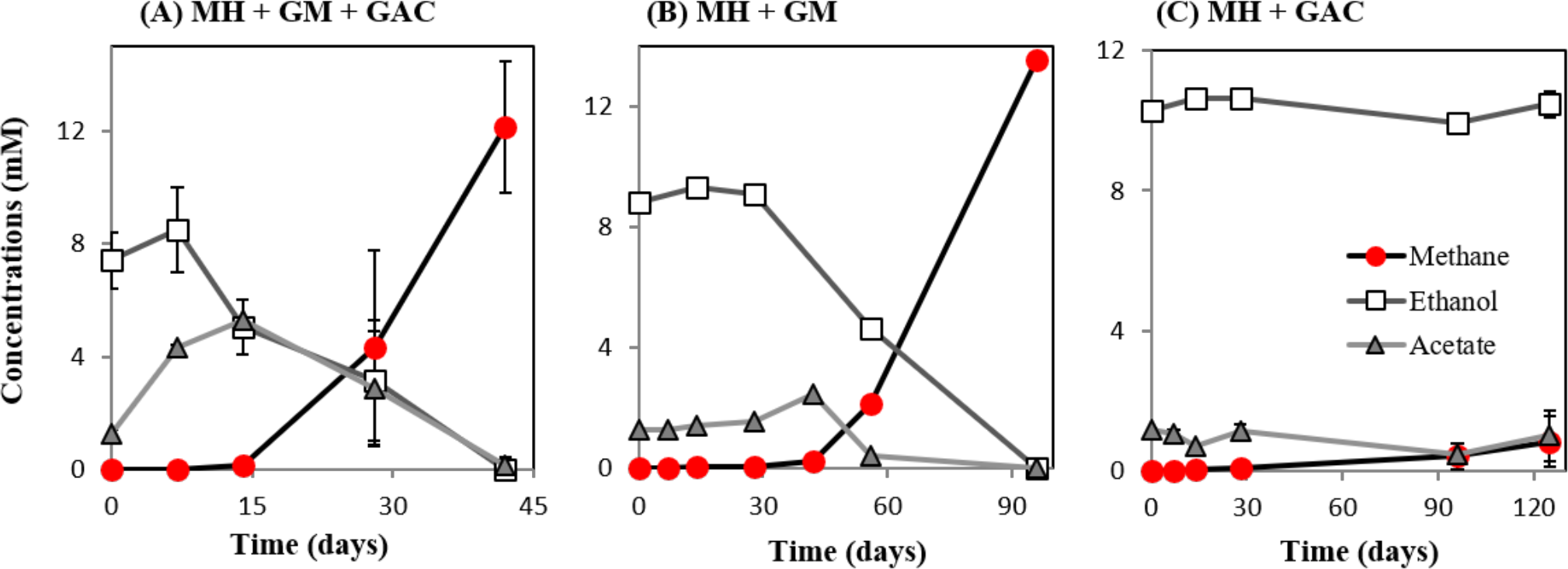
Co-culture experiments with *M. horonobensis* and *G. metallireducens*. *M. horonobensis* established successful co-cultures with G. metallireducens as apparent from ethanol utilization and methane production in the presence (A, n=3) or absence of GAC (B, n=1; see replication in Fig. S6). Alone, *M. horonobensis* could not utilize ethanol or produce methane in the presence of GAC (C, n=3). MH - *Methanosarcina horonobensis*, GM - *Geobacter metallireducens*, GAC - granular activated carbon.

Surprisingly, *M. horonobensis*, which could grow by DIET, was incapable of electromethanogenesis (Fig. 7). Thus we compared the genomes of the two *Methanosarcina, M. horonobensis* and *M. barkeri* to further explain why they were both capable of DIET, but showed dissimilar activities on cathodes at − 400mV.

**Fig. 7.**
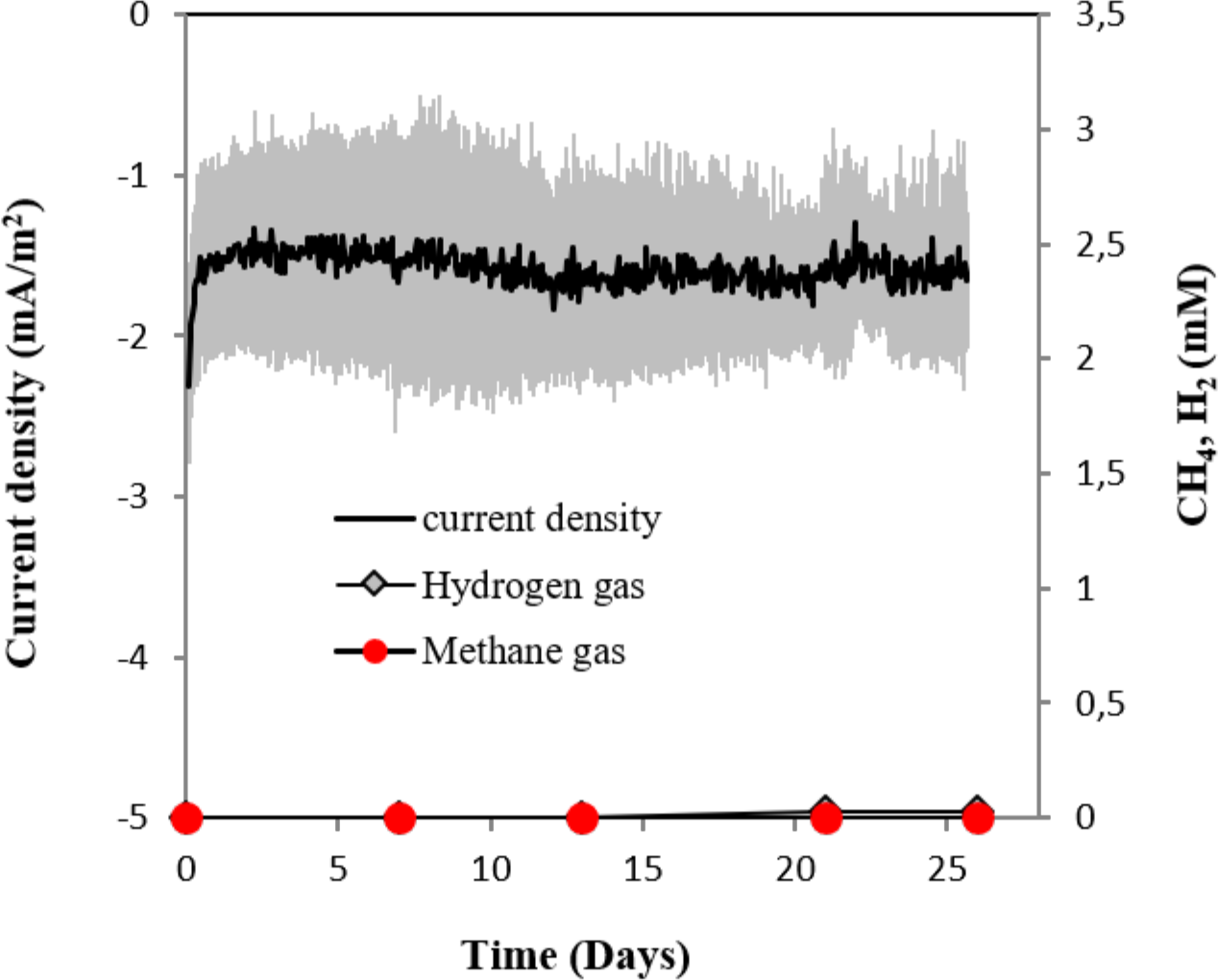
Current consumption and gas production in culturesof *M. horonobensis* (n=3) provided with a cathode poised at − 400 mV (vs. SHE) as sole electron donor.

The main difference between the genomes of *M. barkeri* and *M. horonobensis*, was related to their energy metabolism (Table 1). *M. barkeri* utilizes an energy-converting hydrogenase (Ech) (Kulkarni et al., 2018), which couples the reduction of protons with ferredoxin (Fdx^-^) to the production of a proton motive force according to the reaction: Fdx^−^ (red) + 2H^+^ → Fdx (ox) + H_2_ + ΔμH^+^/ ΔμNa^+^ (Thauer et al., 2008). *M. horonobensis* does not have the Ech (Table 1). An alternative to Ech is the Na^+^-pumping Rnf complex described biochemically in *M. acetivorans* (Schlegel et al., 2012; Suharti et al., 2014), and predicted via genome mining in *M. thermophila* (Wang et al., 2011) and ANME-2 archaea (Wang et al., 2014). Since we could not find any Ech in the genome of *M. horonobensis*, we screened for the genes encoding an Rnf-complex. In *M. horonobensis*, we found all eight representative Rnf-genes (including the cytochrome subunit and Rnf A to G; MSHOH_3554 to 3561), which showed 65-91% protein identity to their *M. acetivorans* counterparts (MA_0658 to 0665).

**Table 1.** Relevant genotypic differences between the methanogens tested during this study

| Species | Energy conservation | S-layer proteins | Predicted c-type cytochromes (CxxCH motif proteins) | Other cytochromes | Predicted Ferredoxins | Predicted thioredoxins |
| --- | --- | --- | --- | --- | --- | --- |
| *Methanosarcina barkeri* MS | Ech-hydrogenase | 8 | 20 (0/1 multiheme*) | 3 (cyt b) | 4 | 10 |
| *Methanosarcina horonobensis* HB-1 | Rnf-complex | 9 | 30 (3 multiheme) | 3 (cyt b) | 6 | 8 |
| *Methanobacterium formicicum* DSM1535 | EhaA/EhbA hydrogenase | None | 16 (None) | None | 4 | 2 |

\*The predicted multiheme cytochrome in *M. barkeri* strain MS had one standard CxxCH and one CxCH motif.

Both Ech and Rnf contain Fe-S centers (Welte and Deppenmeier, 2014), however, the Rnf complex has an accompanying *c*-type cytochrome (Suharti et al., 2014) possibly influencing the overall redox-chemistry on the cell surface. We presume that differences in surface redox chemistry will impact how different *Methanosarcina* interact with extracellular electron donors. Thus, electromethanogenesis at a set potential of – 400 mV is unlikely to match the redox requirements of each type of *Methanosarcina*. On the other hand, in co-cultures, *Geobacter* may coordinate its cytochrome expression to match the redox potential of the partner methanogen, who plays the role of a terminal electron acceptor. This is supported by previous studies showing *Geobacter* modulates their cell-surface proteins to match the electron acceptor provided (Ishii et al., 2018; Otero et al., 2018).

When contrasting the genomes of the two *Methanosarcina* species we also observed significant differences regarding nitrogen fixation, mobile elements, and sensing/chemotaxis proteins (Table 2). As such, compared to *M. horonobensis, M. barkeri* encodes for more N_2_-fixation proteins (86%). Compared to *M. barkeri, M. horonobensis* encodes for more small-molecule-interaction proteins such as redox-sensing and chemotaxis proteins (185%) and mobile elements than *M. barkeri* (16 fold increase) (Table 2). The exact role of these proteins in extracellular electron uptake by these *Methanosarcinas* is unknown and warrants further investigation.

**Table 2.**
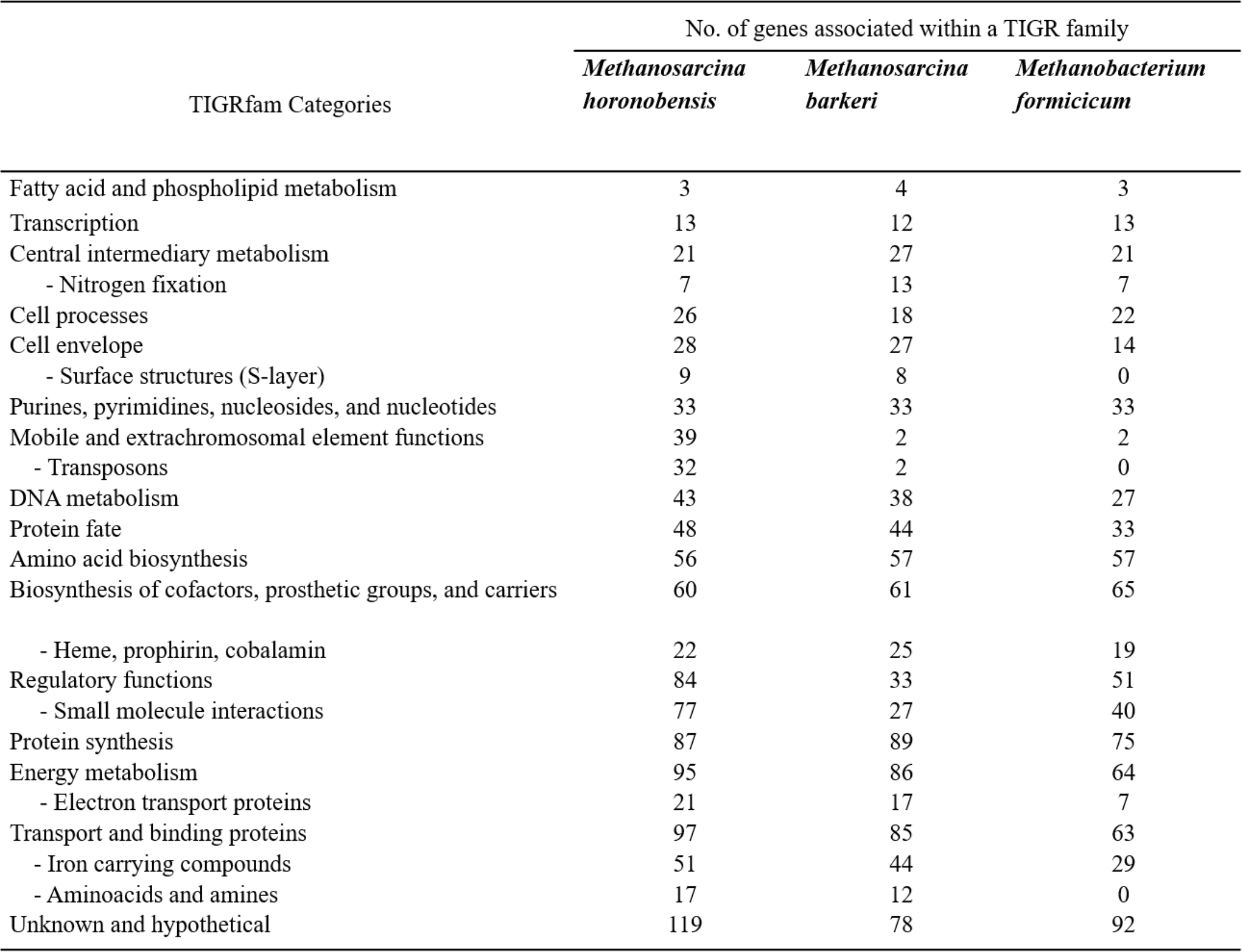
Genomic comparison of three methanogens based on TIGR family protein categories.

Furthermore, to determine why *Methanosarcina* could do DIET, but not *Methanobacterium*, we compared the genomes of the two *Methanosarcina* species with that of *M. formicicum* (Table 2). In contrast to the *Methanobacterium*, both *Methanosarcina* species encode in their genomes three times the amount of genes for electron transport proteins and circa 50% more genes for cell surface and transport proteins (Table 2). Especially, outer surface S-layer proteins were only present in the two *Methanosarcina* (Table 2). S-layer proteins were previously suggested to play a role in extracellular electron transfer in *Methanosarcina* related ANME-2, which carry anaerobic methane oxidation syntrophically (McGlynn, 2017; McGlynn et al., 2015; Timmers et al., 2017). Future gene-expression and deletion studies could shed light on the possible role of S-layer proteins in DIET-interactions.

## Conclusion

Three methanogens were investigated for their ability to do extracellular electron uptake from (1) a cathode at −400 mV, (2) directly from an electrogenic-DIET partner or (3) from a DIET-partner, but mediated by conductive particles. Only *M. barkeri* was able to carry out all three forms of extracellular electron uptake, making this the first observation of a *Methanosarcina* in pure culture performing electromethanogenesis. The conditions in our abiotic electrochemical controls did not lead to H_2_-evolution at −400mV, according to electrochemical and analytical tests. Therefore, under these conditions, it was impossible to sustain a methanogen with high H_2_-affinity, like *M. formicicum.* Besides *M. formicicum* was incapable to retrieve electrons directly from the electrode or from a DIET partner (direct or via conductive particles). In this study, we also demonstrated that another *Methanosarcina*, *M. hornobensis* performed DIET with *Geobacter* (direct or via conductive particles). However, surprisingly, *M. horonobensis* was incapable of electromethanogenesis. We screened the genomes of the two *Methanosarcina* and identified differences (e.g. energy metabolism), which could lead to phenotypic variability and thus contrasting electromethanogenesis-ability. Compared to *M. barkeri*, *M. horonobensis* is a better candidate for understanding electron uptake from a DIET syntrophic partner. This is because unlike *M. barkeri, M. horonobensis* does not utilize H_2_, and it grows as single cells on freshwater media, which is ideal for genetic studies.

## Acknowledgments

The Innovationsfond grant number 4106-00017 funded this work. We would like to thank Lasse Ørum Smidt and Heidi Grøn Jensen for lab assistance.

## Author contribution

MY and A-ER conceived the study with support from BT and LO. MY performed all experiments with support from OS. MY analyzed the data with support from A-ER. MY wrote the manuscript with help from A-ER. All authors (MY, A-ER, BT, OS and LO) contributed to drafting and editing the manuscript.

