## Supplementary File 1 for "Extracellular Electron Uptake by Two *Methanosarcina* Species"

**Supplementary Figures**

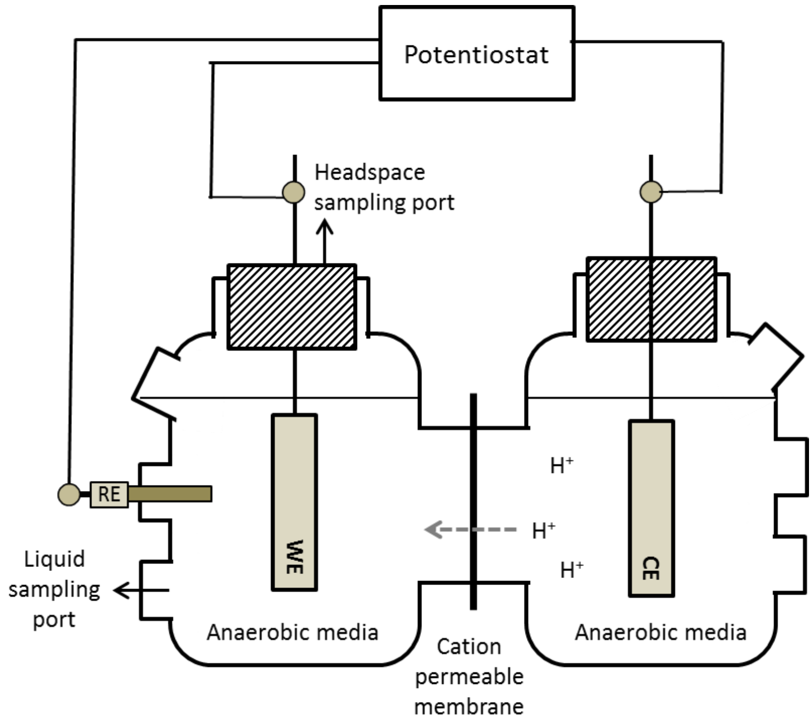


**(Fig. S1)** Two chambered electrochemical reactors separated by a proton exchange membrane. RE; reference electrode, WE; working electrode, CE; counter electrode.

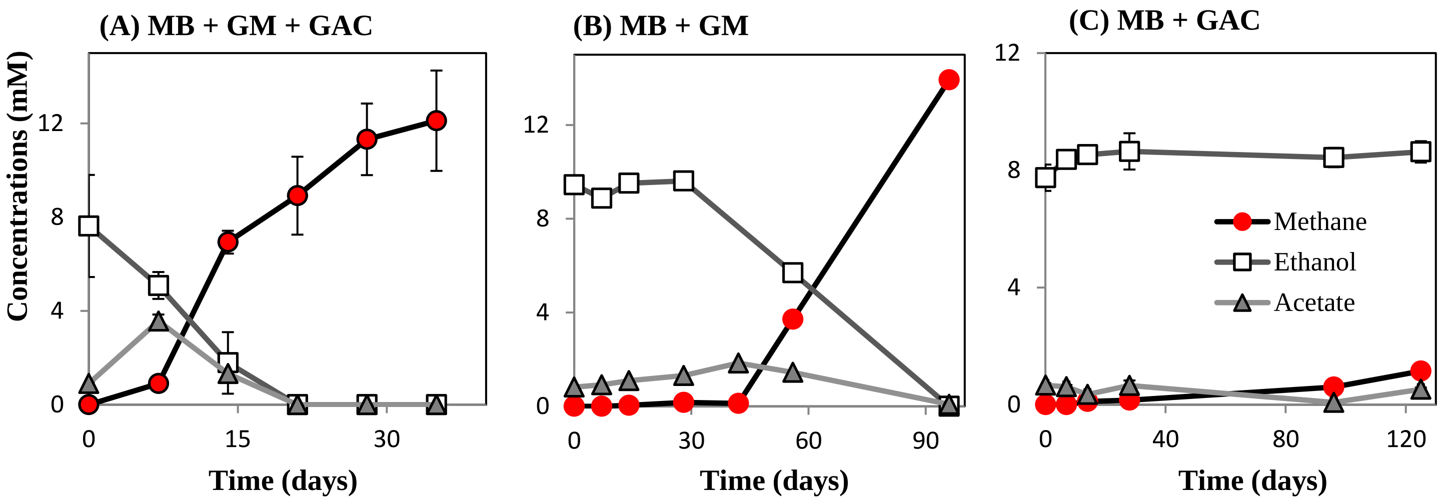


**(Fig. S2)** Co-cultures of *M. barkeri* with *G.metallireducens* feeding on ethanol with or without GAC. MH; *Methanosarcina barkeri*, GM; *Geobacter metallireducens*, GAC; granular activated carbon, Circles; Methane, Squares; Ethanol, Triangles; Acetate. The results are from triplicate samples except for (B) which is a representative sample for which the rest of the replicate data is presented in Fig S6.

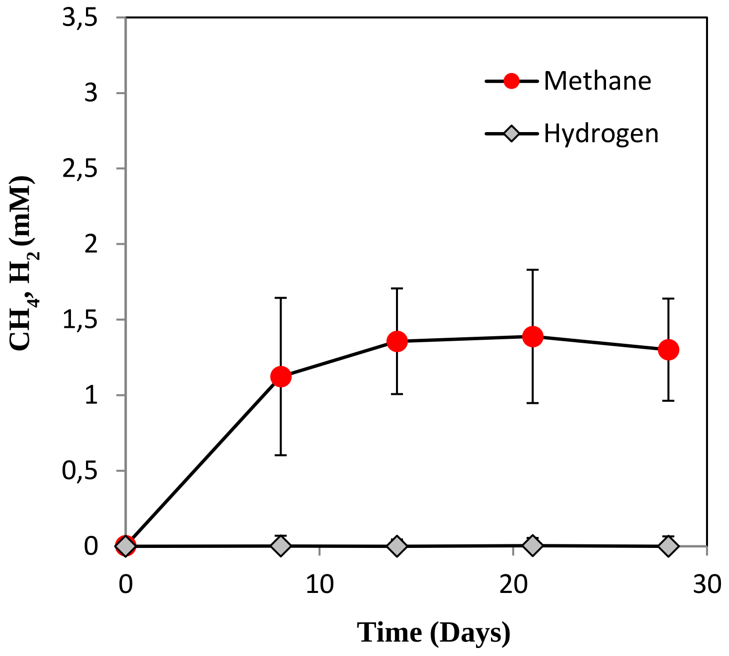


**(Fig. S3)** Open circuit control reactors *M. barkeri*. Residual methane production from substrate carry-over introduced together with the inoculum. Error bars are based on standard deviation (n=3).

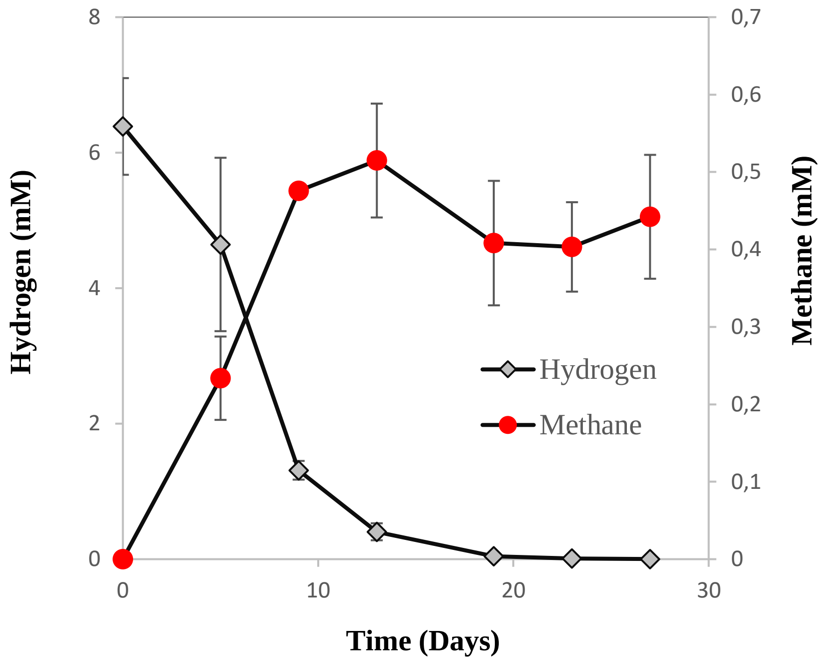


**(Fig. S4)** Hydrogen consumption and methane production from *M. formicicum* reactors at -400 mV (vs. SHE) with added Hydrogen gas in the headspace. Error bars are based on standard deviation (n=2).

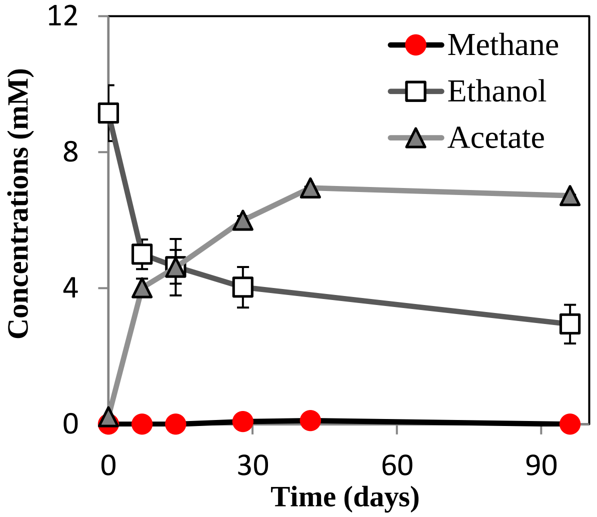


**(Fig. S5)** *G.metallireducens* with ethanol as the sole substrate ammended with granular activated carbon. Error bars are based on standard deviation (n=3). Circles; Methane, Squares; Ethanol, Triangles; Acetate .

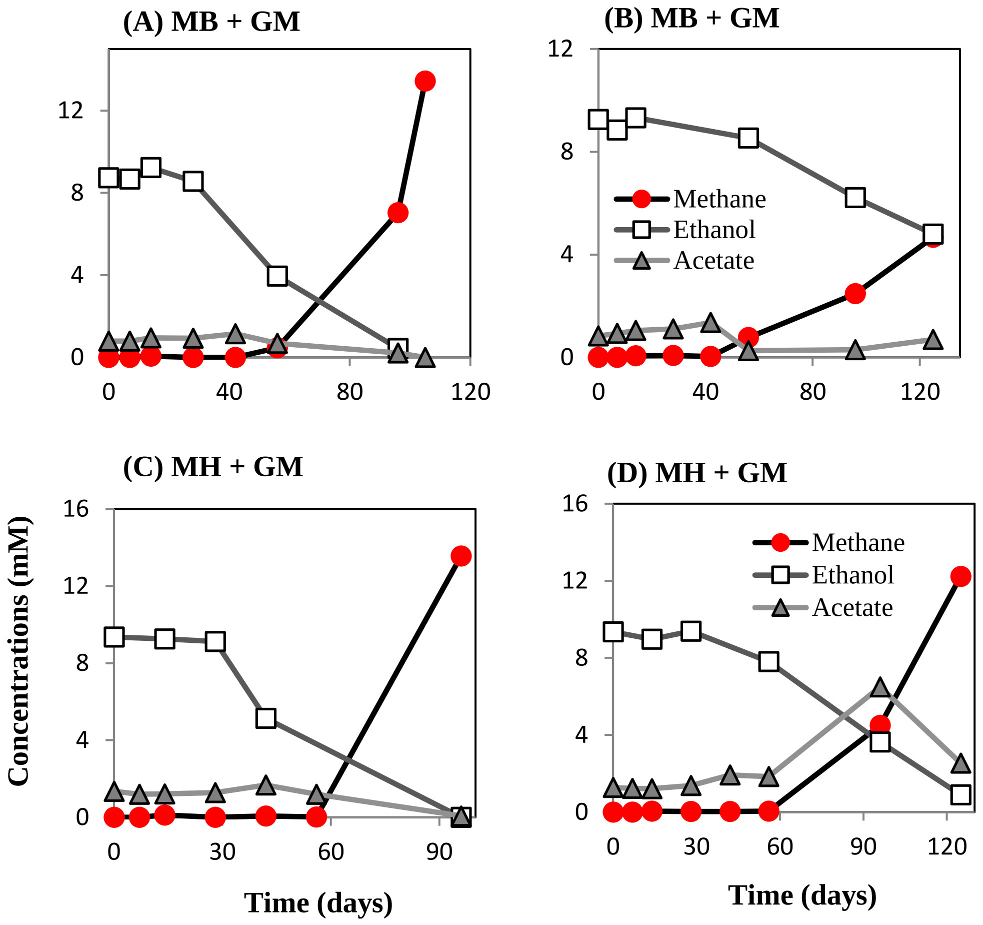


**(Fig. S6)** Replicate co-cultures of *Methanosarcina* sp. with *G. metallireducens* (A to D) feeding on syntrophically oxidizing ethanol without the addition of external electron shuttles. MB; *Methanosarcina barkeri,* MH*; Methanosarcina horonobensis,* GM*; Geobacter metallireducens,* Circles; Methane, Squares; Ethanol, Triangles; Acetate .
